# Detrimental Impact of a Type VI Secretion System on Direct Interspecies Electron Transfer

**DOI:** 10.1101/2023.03.02.530876

**Authors:** Jessica A. Smith, Dawn E. Holmes, Trevor L. Woodard, Yang Li, Xinying Liu, Li-Ying Wang, David Meier, Ingrid A. Schwarz, Derek R. Lovley

**Author notes:** These authors contributed equally to this work (Jessica A. Smith, Dawn E. Holmes).

## Abstract

Direct interspecies electron transfer (DIET) is important in anaerobic communities of environmental and practical significance. Other than the need for close physical contact for electrical connections, the interactions of DIET partners are poorly understood. Type VI secretion systems (T6SSs) typically kill competitive microbes. Surprisingly, *Geobacter metallireducens* highly expressed T6SS genes when DIET-based co-cultures were initiated with *Geobacter sulfurreducens*. T6SS gene expression was lower when the electron shuttle anthraquinone-2,6-disulfonate was added to alleviate the need for interspecies contact. Disruption of *hcp*, the *G. metallireducens* gene for the main T6SS needle-tube protein subunit, and the most highly upregulated gene in DIET-grown cells, eliminated the long lag periods required for the initiation of DIET. The mutation did not aid DIET in the presence of granular activated carbon, consistent with the fact that DIET partners do not make physical contact when electrically connected through conductive materials. The *hcp*-deficient mutant also established DIET quicker with *Methanosarcina barkeri*. However, the mutant also reduced Fe(III) oxide faster than the wild-type strain, a phenotype not expected from the loss of the T6SS. Quantitative PCR revealed greater gene transcript abundance for key components of extracellular electron transfer in the *hcp*-deficient mutant versus the wild-type strain, potentially accounting for the faster Fe(III) oxide reduction and impact on DIET. The results highlight that interspecies interactions beyond electrical connections may influence DIET effectiveness. The unexpected increase in the expression of genes for extracellular electron transport components when *hcp* was deleted emphasize the complexities in evaluating the electromicrobiology of highly adaptable *Geobacter* species.

**IMPORTANCE:** Direct interspecies electron transfer (DIET) is an alternative to the much more intensively studied process of interspecies H_2_ transfer as a mechanism for microbes to share electrons during the cooperative metabolism of energy sources. DIET is an important process in anaerobic soils and sediments generating methane, a significant greenhouse gas. Facilitating DIET can accelerate and stabilize the conversion of organic wastes to methane biofuel in anaerobic digesters. Therefore, a better understanding of the factors controlling how fast DIET partnerships are established is expected to lead to new strategies for promoting this bioenergy process. The finding that when co-cultured with *G. sulfurreducens, G. metallireducens* initially expressed a type VI secretion system, a behavior not conducive to interspecies cooperation, illustrates the complexity in establishing syntrophic relationships.

## INTRODUCTION

A better understanding of the physiological characteristics of microbes that participate in direct interspecies electron transfer (DIET) is required in order to determine how both natural and engineered anoxic environments function (1-3). For example, DIET appears to be the primary route for electron exchange between electron-donating bacteria and electron-accepting partners in some types of anaerobic digesters (4, 5). In digesters in which interspecies H_2_ transfer predominates, modifying operating conditions to enhance DIET can accelerate and stabilize the conversion of organic wastes to methane, a needed improvement to this important bioenergy strategy (3, 6). Molecular studies have demonstrated that DIET may be a major process in terrestrial methanogenic environments that are significant sources of atmospheric methane (7), a conclusion that is further supported by the reinterpretation of data on H_2_ fluxes in these environments (8).

Most of the initial research following the discovery of DIET (9) focused on identifying which microbes have the potential to participate in DIET and the organic substrates that can support DIET (5, 8, 10-21). Study of the expression of genes and proteins that enhance electron exchange between species has also been emphasized (5, 9, 10, 13, 22-25). However, other adaptations that promote the switch from a free-living existence to living in close physical association, as is necessary to establish electrical connections for DIET, seem likely.

The expression of type VI secretion systems (T6SSs) is expected to be antithetical to interspecies cooperation. Approximately 25% of Gram-negative bacteria have T6SSs that form contractile nanomachines that inject toxins directly into other microbes to eliminate their competition (26-36). T6SSs are important in such polymicrobial environments as the human colon (37-41), cow rumen (42, 43), the plant rhizosphere (44, 45), the light organ of the bobtail squid (46), and soil (47-49). In some instances, T6SSs can also be involved in such non-antagonistic behaviors as the modulation of quorum sensing and stress response (50), self-recognition (51-53), and the acquisition of various metals such as zinc, copper, manganese, or iron (48-51, 54-57).

Molecular analyses have demonstrated that *Geobacter* species are important electron-donating partners for DIET in natural environments, such as subsurface terrestrial soils (7) as well as in some anaerobic digesters (4, 5). The often-observed enrichment of *Geobacter* when methane production is stimulated with the addition of conductive materials provides further circumstantial evidence for a role of *Geobacter* in DIET (3, 6, 58). The availability of pure cultures of genetically tractable *Geobacter* species that can participate in DIET in defined co-cultures has enabled elucidation of important electrical contacts for DIET, such as multi-heme *c*-type cytochromes and electrically conductive pili (5, 9, 10, 23, 25, 59), as well as strategies for enhancing DIET with electrically conductive minerals and carbon materials (60-64).

However, *Geobacter* species are also often found to be free-living in anaerobic soils and sediments, typically transferring electrons to extracellular electron acceptors such as Fe(III) oxides and humic substances (65). T6SS genes are present in some but not all *Geobacter* genomes (66, 67) (Table S1; Fig. S1). It might be expected that T6SSs could be beneficial to free-living *Geobacter* species competing against other microbes for resources, but not for developing syntrophic cooperation. Here we report that *Geobacter metallireducens* highly expresses genes coding for its T6SS in the initial stages of establishing a DIET-based co-culture with *Geobacter sulfurreducens*, a factor possibly lengthening the adaptation period required for DIET-based growth of the co-culture and accounting for the ability of conductive materials to accelerate DIET.

## Materials and Methods

### Laboratory strains and culture conditions

*Geobacter* cultures were obtained from our laboratory culture collection and routinely cultured under strict anaerobic conditions, as previously described (68). *G. metallireducens* was grown in Fe(III) citrate (FC) medium (69) with 20 mM ethanol provided as the sole electron donor and 56 mM Fe(III) citrate as the sole electron acceptor, or with 20 mM acetate as the donor and 50 mM Fe(III) oxide as the acceptor. *G. sulfurreducens* was grown in medium with 10 mM acetate provided as the sole electron donor and 40 mM fumarate as the sole electron acceptor (NBAF medium) (68). Co-cultures were initiated with equal amounts of both organisms in anaerobic pressure tubes containing 10 mL of NBF medium (acetate-free NBAF), with 10 mM ethanol provided as the sole electron donor and 40 mM fumarate as the electron acceptor. When noted, additions of anthraquinone-2,6, -disulfonate (AQDS) were made from a concentrated stock to provide a final concentration of 50 μM. In some instances, granular activated carbon (GAC; 8-20 mesh (Sigma-Aldrich)) was added at 0.1g/ 10 ml.

*Methanosarcina barkeri* was obtained from our laboratory culture collection and grown in co-culture with *G. metallireducens* with ethanol as the electron donor, as previously described (59).

### Analytical techniques

Organic acids were monitored with high performance liquid chromatography (HPLC), as previously described (70). Changes in ethanol concentration were monitored with gas chromatography, as previously described (4). Methane was monitored in the headspace by gas chromatography with a flame ionization detector (SHIMADZU, GC-8A), as previously described (71). Fe(II) concentrations were determined by first incubating samples for 1 hour in 0.5N HCl and then measuring Fe(II) with a ferrozine assay at an absorbance of 562 nm (72).

### Illumina sequencing and data analysis

For all experimental conditions, total RNA was extracted from triplicate samples at mid-log phase growth when succinate concentrations reached approximately 25 mM using the RNeasy Mini Kit (Qiagen) according to the manufacturer’s instructions. Samples were treated with Turbo DNA-free DNase (Ambion, Austin, TX), and the RNA samples were tested for genomic DNA (gDNA) contamination by PCR amplification of the 16S rRNA gene. mRNA was enriched using the MICROB*Express* kit (Ambion), according to the manufacturer’s instructions.

Directional libraries were prepared with the ScriptSeq™ v2 RNA-Seq Library Preparation Kit (Epicentre) and single end sequencing was performed on a Hi-Seq 2000 platform at the Deep Sequencing Core Facility at the University of Massachusetts Medical School in Worcester, Massachusetts. The program FASTQC (http://www.bioinformatics.babraham.ac.uk/projects/fastqc/) was used to visualize and quality check all raw data. Initial raw non-filtered libraries contained an average of 13175155.5 +/-1758892 and 10227370.2 +/ 1558219.6 reads in the DIET and quinone-mediated interspecies electron transfer (QUIET) libraries that were ∼100 base pairs long. Sequences from all of these libraries were trimmed and filtered with Trimmomatic (bolger 2014) yielding an average of 9286241 +/-1665081.9 and 12393095 +/-1719373.9 reads for the DIET and QUIET libraries.

### Mapping of mRNA reads

Trimmed and filtered mRNA reads from triplicate samples for the two different co-culture conditions (DIET and QUIET) were mapped against the *G. metallireducens* strain GS-15 genome (NC_007517) and the *G. sulfurreducens* strain PCA genome (NC_002939) downloaded from IMG/MER (img.jgi.doe.gov) using ArrayStar software (DNAStar). Common dispersion (Disp) and biological coefficient of variation (BCV) values between DIET and QUIET replicates were calculated with the edgeR package in Bioconductor (73). Common dispersion and BCV values for DIET and QUIET libraries were Disp=0.01953 and BCV=0.1398 and Disp=0.11752 and BCV=0.3428, respectively. A multidimensional scaling (MDS) plot was also generated with edgeR software and showed that replicates from the DIET and QUIET libraries clustered together but separately from each other (Fig. S2).

Once the quality of RNAseq libraries was determined, differential expression studies were done with the edgeR package in Bioconductor (73). Genes with p-values ≤ 0.05 and fold changes ≥ 2 were considered differentially expressed. Using these criteria, 945 *G. metallireducens* genes and 967 *G. sulfurreducens* genes were up-regulated in DIET-grown co-cultures and 603 *G. metallireducens* genes and 848 *G. sulfurreducens* genes were up-regulated in QUIET-grown co-cultures (Table S2).

### Quantitative RT-PCR

Quantitative RT-PCR was conducted with mRNA extracted from triplicate cultures of *G. metallireducens* wild-type and Δ*hcp* (Gmet_0280) strains grown by Fe(III) oxide respiration, in co-culture with *M. barkeri*, or in co-culture with *G. sulfurreducens*. Cells were harvested during the mid-logarithmic phase by centrifugation at 4,000 rpm for 15 min at 4°C. After centrifugation, the pellets were frozen in liquid nitrogen and stored at -80°C until RNA extraction procedures were performed. Total RNA from sample pellets was extracted as previously described (74). Complementary DNA (cDNA) was generated from mRNA using the Invitrogen SuperScript IV First Strand Synthesis System (ThermoFisher Sci).

Primer pairs used for qRT-PCR are provided in Table S3. Three different housekeeping genes were used as external controls; *recA* which codes for recombinase A, *proC* which codes for pyrroline-5-carboxylate reductase, and *rpoB* which codes for the beta subunit of RNA polymerase. Power SYBR green PCR master mix (Applied Biosystems, Foster City, CA) and an ABI 7500 real-time PCR system were used to amplify and to quantify all PCR products. Each reaction mixture consisted of forward and reverse primers at a final concentration of 200 nM, 5 ng of gDNA, and 12.5 μl of Power SYBR green PCR master mix (Applied Biosystems). Relative levels of expression of the studied genes were calculated by the 2−^ΔΔ*CT*^ threshold cycle (CT) method (75).

### Mutant construction

Primers used for construction of gene replacement mutants and complement strains are listed in Table S4. Deletion mutants were made by replacing the gene of interest with a spectinomycin antibiotic resistance cassette (76). All restriction digestions were carried out according to manufacturer’s instructions. PCRs were done using the JumpStart Taq DNA polymerase (Sigma-Aldrich). Primer pairs were used to amplify by PCR flanking regions of approximately 500 bp downstream and upstream of the target genes using the appropriate genomic DNA as a template. PCR products were digested with the *AvrII* (CCTAGG) (NEB, Beverly, MA) restriction endonuclease, ethanol precipitated, and ligated with T4 DNA ligase (NEB). The ligation reaction was loaded onto a 1% agarose gel, and a 1 kb band was purified using the Qiaquick Gel Extraction Kit (Qiagen) and cloned into pCR2.1 TOPO cloning vector. Sequences of the cloned products were verified by Sanger sequencing. The spectinomycin cassette was digested with *XbaI* (TCTAGA) (NEB) from pUC19*-Spr*^*r*^ *loxP* (76), and the recombinant plasmid was digested with *AvrII*. The spectinomycin resistance cassette was cloned into the plasmid to complete the construction of the mutant alleles. Plasmids bearing mutant alleles were linearized and concentrated by ethanol precipitation. The linearized plasmids were electroporated as described previously (76). Antibiotics were added for selection purposes only. Replacement of wild type alleles by mutant alleles was verified by PCR and Sanger sequencing.

Gmet_0280 was complemented *in trans* by amplifying the gene with its native ribosome binding site (RBS) using *G. metallireducens* genomic DNA as a template. The resulting PCR product was then digested and cloned under control of a constitutive lac promoter into pCM66 (77) and electroporated into the Gmet_0280-deficient strain, as previously described (76).

### Data availability

Illumina sequence reads have been submitted to the SRA NCBI database under BioProject PRJNA722959 and Biosample SAMN18796025.

## Results and discussion

### Increased expression of *G. metallireducens* T6SS genes during DIET

Comparing gene expression patterns in defined co-cultures growing via DIET versus co-cultures growing with the exchange of diffusible electron shuttles has proven to be an effective strategy for identifying mechanisms for electron transfer during DIET (23, 24). Therefore, co-cultures of *G. metallireducens* and *G. sulfurreducens* were grown in medium with ethanol as the electron donor and fumarate as the electron acceptor as previously described (9, 78). The two species must cooperate to share electrons in this medium because only *G. metallireducens* can utilize ethanol as an electron donor and only *G. sulfurreducens* uses fumarate as electron acceptor (9). AQDS (50 µM) was added to one set of co-cultures to promote quinone-mediated interspecies electron transfer (QUIET) in which the two species remain free-living and AQDS serves as an electron shuttle between them (78). However, in the absence of AQDS the two species must form tight physical association for direct electron transfer from *G. metallireducens* to *G. sulfurreducens* because *G. metallireducens* cannot produce H_2_ or formate as electrons shuttles when metabolizing ethanol (22, 23).

The initial AQDS-amended and unamended co-cultures were sampled for gene expression analysis after they had reduced ca. 25 mM of the 40 mM fumarate available. DIET-grown *G. sulfurreducens* had higher transcript abundances for genes previously found to be important in DIET (Table S2c). These included genes for over 25 *c*-type cytochromes, including *omcS, omcB, omcX*, and *omcI*, which encode multi-heme, outer-surface *c*-type cytochromes. The gene coding for the pilin monomer, PilA, which is assembled into electrically conductive pili (79) was also 63-times more highly expressed in DIET-grown cells.

Transcripts for 19 different *c*-type cytochrome genes were also ≥2-fold more abundant in DIET-grown *G. metallireducens* (Table S2a). These included Gmet_0930, which codes for an octaheme outer membrane *c*-type cytochrome and Gmet_0910, the gene for the outer membrane *c*-type cytochrome, OmcF, from the PccF porin-cytochrome complex (59). Both Gmet_0930 and Gmet_0910 are important for Fe(III) oxide reduction and DIET-based growth (17, 59, 80). Gmet_2029, which codes for a lipopolysaccharide protein likely to be involved in biofilm formation and required for Fe(III) oxide reduction (80), was expressed more than 3 fold higher in DIET-grown cells, but *pilA* was not more highly expressed by DIET-grown *G. metallireducens* cells (Table S2a).

Other genes that would not be expected to be involved in electron transfer were also more highly expressed in *G. metallireducens* cells growing by DIET (Table 1; Table S2a). The two genes with the greatest increase in abundance of transcripts in DIET-versus QUIET-grown cells were Gmet_2080 and Gmet_2078, annotated as ‘T6SS needle tube protein TssD’ and ‘T6SS protein ImpB’. Transcripts for other T6SS proteins were also much more abundant in DIET-versus QUIET-grown *G. metallireducens* (Table 1). This included 13 of the genes needed to construct the T6SS nanomachine in other microbes (26, 31, 81). Furthermore, genes coding for all putative T6SS effectors, immunity proteins, and effector chaperones in *G. metallireducens*, with the exception of Gmet_0291 which codes for a putative chaperone protein, were at least 5-times more highly expressed in DIET-grown *G. metallireducens* cells (Table 1). Only some of the *G. sulfurreducens* T6SS-related genes were highly expressed in DIET-versus QUIET-grown cells (Table S5).

**Table 1.** Differences in transcript abundance for all 13 core T6SS genes in *G. metallireducens* grown via DIET compared to QUIET and for genes coding for putative T6SS effector and associated immunity proteins (P-value cutoff ≤ 0.05). ND: no difference tse: T6SS effector protein; tsi: T6SS immunity protein

| Gene | Name | Annotation | Fold-change<br>DIET vs.<br>QUIET | p-value |
| --- | --- | --- | --- | --- |
| **Core T6SS Genes** |  |  |  |  |
| Gmet_0273 | *tssL* | membrane core complex protein | 129.56 | $1.38 \times 10^{-6}$ |
| Gmet_0274 | *tssM* | membrane core complex protein | 47.98 | $8.99 \times 10^{-26}$ |
| Gmet_0275 | *tssA* | baseplate complex protein | 91.34 | $2.71 \times 10^{-23}$ |
| Gmet_0278 | *tssB* | Sheath protein | 703.55 | $1.71 \times 10^{-21}$ |
| Gmet_0279 | *tssC* | Sheath protein | 44.18 | $4.29 \times 10^{-27}$ |
| Gmet_0280 | *hcp* | inner tube protein | 1397.32 | $1.74 \times 10^{-30}$ |
| Gmet_0286 | *vrgG* | puncturing device | 44.88 | $9.12 \times 10^{-15}$ |
| Gmet_3310 | *tssJ* | membrane core complex protein | 27.08 | $5.50 \times 10^{-10}$ |
| Gmet_3311 | *tssK* | baseplate complex protein | 5.69 | $4.86 \times 10^{-8}$ |
| Gmet_3312 | *tssE* | baseplate complex protein | 44.53 | $3.93 \times 10^{-3}$ |
| Gmet_3313 | *tssF* | baseplate complex protein | 136.45 | $7.68 \times 10^{-7}$ |
| Gmet_3314 | *tssG* | baseplate complex protein | 52.36 | $1.78 \times 10^{-3}$ |
| Gmet_3315 | *tssH* | ClpV1 protease involved in sheath recycling | 11.47 | $5.50 \times 10^{-11}$ |
| **Effector/ Immunity Genes** |  |  |  |  |
| Gmet_0284 | *tsi1* | NTF-domain protein (immunity protein) | 14.01 | $1.02 \times 10^{-8}$ |
| Gmet_0285 | *tse1* | peptidoglycan-binding D-alanyl-D-alanine<br>carboxypeptidase | 15.92 | $9.05 \times 10^{-10}$ |
| Gmet_0287 | *tse2* | fatty acid metabolism protein | 5.16 | $7.36 \times 10^{-5}$ |
| Gmet_0288 | | PAAR-like DUF4150 domain protein | 126.78 | $1.76 \times 10^{-6}$ |
| Gmet_0290 | *tse3* | PGAP1 domain protein; phospholipase | 7.71 | $1.31 \times 10^{-8}$ |
| Gmet_0291 |  | DUF2169 domain chaperone protein | ND |  |

### Disrupting the *hcp* gene in *G. metallireducens* accelerates adaption to DIET

The high expression of T6SS genes in *G. metallireducens* during growth via DIET was surprising because a primary function of the T6SS is elimination of competing species (31, 32, 34, 82-84). To determine whether expression of the T6SS by *G. metallireducens* impacted DIET, the gene for the Hcp needle-tube protein (Gmet_0280), the most highly differentially expressed gene in DIET-versus QUIET-grown cells (Table S2A), was disrupted by replacing the gene with a spectinomycin resistance cassette.

As previously described (9), co-cultures established in ethanol-fumarate medium with wild-type *G. metallireducens* required over 25 days to begin DIET, monitored as the accumulation of succinate from fumarate reduction (Fig. 1a). In contrast, there was a shorter lag in adaption to DIET in co-cultures initiated with the Hcp-deficient strain of *G. metallireducens* (Fig. 1a). Large aggregates (1-2 mm diameter) were visibly apparent in the co-cultures with the Hcp-deficient strain of *G. metallireducens* even when the co-cultures were first established. In contrast, as previously reported (9), in co-cultures established with wild-type *G. metallireducens* large aggregates only appeared after multiple successive transfers of the co-cultures. The Hcp-deficient strain also produced visible aggregations in Fe(III) citrate medium, which was not observed in the wild-type strain (Fig. S3).

**Figure 1.**
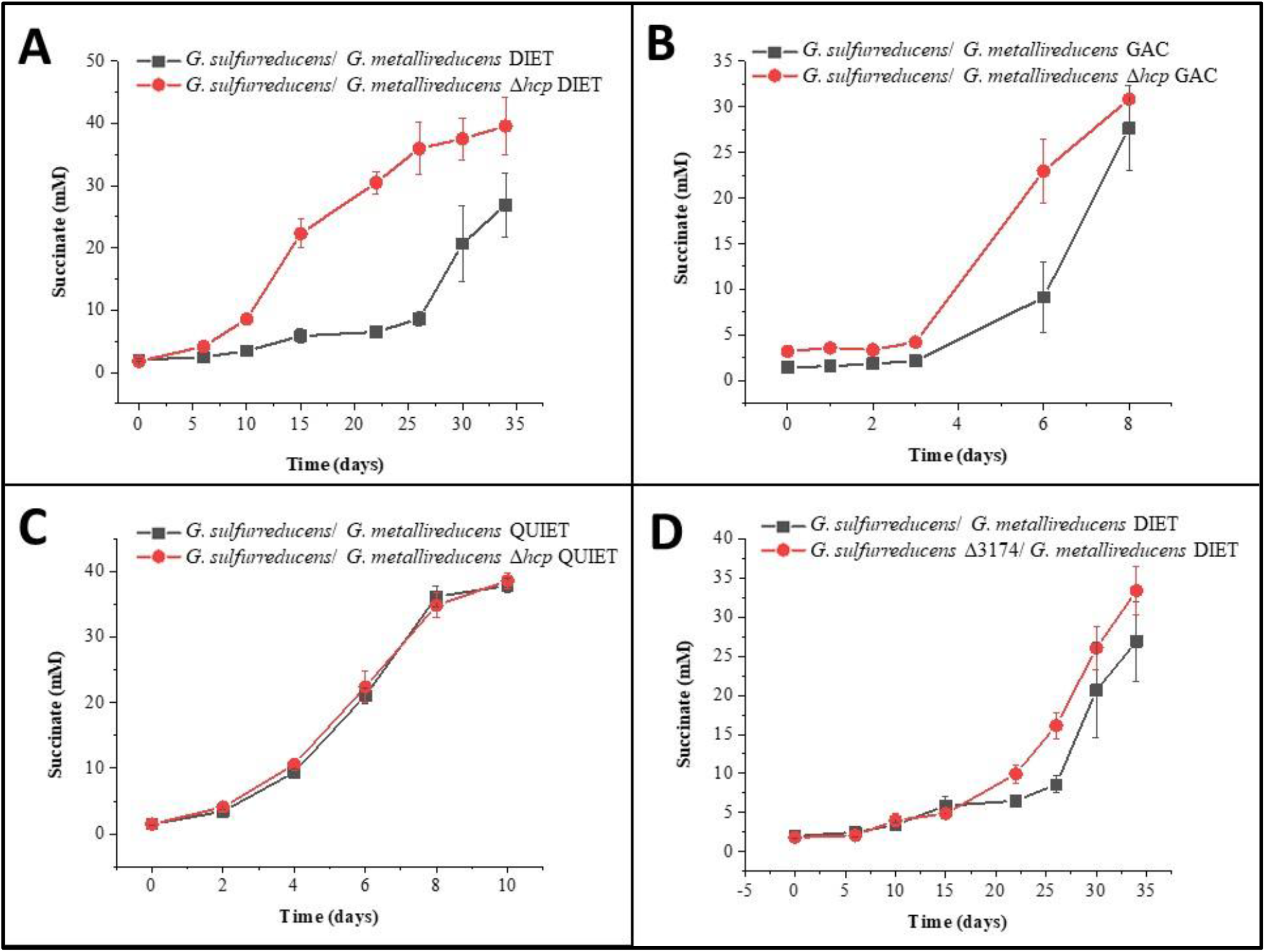
The impact of deletion of a key T6SS component on DIET. Metabolism of (A) DIET co-cultures initiated with the wild-type or *hcp*-deficient strain of *G. metallireducens* and wild-type *G. sulfurreducens*, (B) GAC-supplemented co-cultures initiated with the wild-type or *hcp*-deficient strain of *G. metallireducens* and wild-type *G. sulfurreducens*, (C) QUIET co-cultures initiated with the wild-type or *hcp*-deficient strain of *G. metallireducens* and wild-type *G. sulfurreducens*, and (D) DIET co-cultures initiated with the wild-type or GSU3174-deficient strain of *G. sulfurreducens* and wild-type *G. metallireducens*. Each point and error bars represent the average and standard deviation of triplicate measurements.

The Hcp-deficient *G. metallireducens* strain did not have a substantial advantage over wild-type cells if granular activated carbon (GAC) was added to the co-cultures (Fig. 1b). In the presence of GAC, which is electrically conductive, *G. metallireducens* and *G. sulfurreducens* attach to the GAC surface rather than producing dual-species aggregates and the cells are not close enough for DIET via electrically conductive pili or *c*-type cytochromes (60). Deletion of the genes for these biological electrical connections does not inhibit DIET in the presence of GAC, suggesting that GAC is a highly effective conduit for DIET (60). Co-cultures established with the Hcp-deficient *G. metallireducens* or wild-type *G. metallireducens* grew at the same rate when AQDS was provided as an electron shuttle for QUIET (Fig. 1c). Like GAC, AQDS also eliminates the need for direct cell-to-cell contacts for interspecies electron transfer (78).

Disrupting the gene for Hcp from *G. sulfurreducens* (ΔGSU3174) did not substantially decrease the lag time required for initiation of co-culture metabolism under conditions that require DIET for growth (Fig. 1d). This is consistent with the observation that *G. sulfurreducens* did not increase expression of genes for most T6SS components in DIET-versus QUIET-grown cells (Table S5).

To determine the potential impact of T6SS expression on *Geobacter* interactions with methanogens, co-cultures of *G. metallireducens* and *M. barkeri* were initiated as previously described (10) in medium with ethanol as the electron donor. As previously described (10), there was a lag period of more than 30 days in co-cultures initiated with wild-type *G. metallireducens* (Fig. 2). In contrast, there was very little lag in co-cultures initiated with the Hcp-deficient *G. metallireducens* strain. While studies have focused on T6SSs targeting bacterial and eukaryotic cells (85, 86), the effect of T6SSs on archaeal cells requires further study (87).

**Figure 2.**
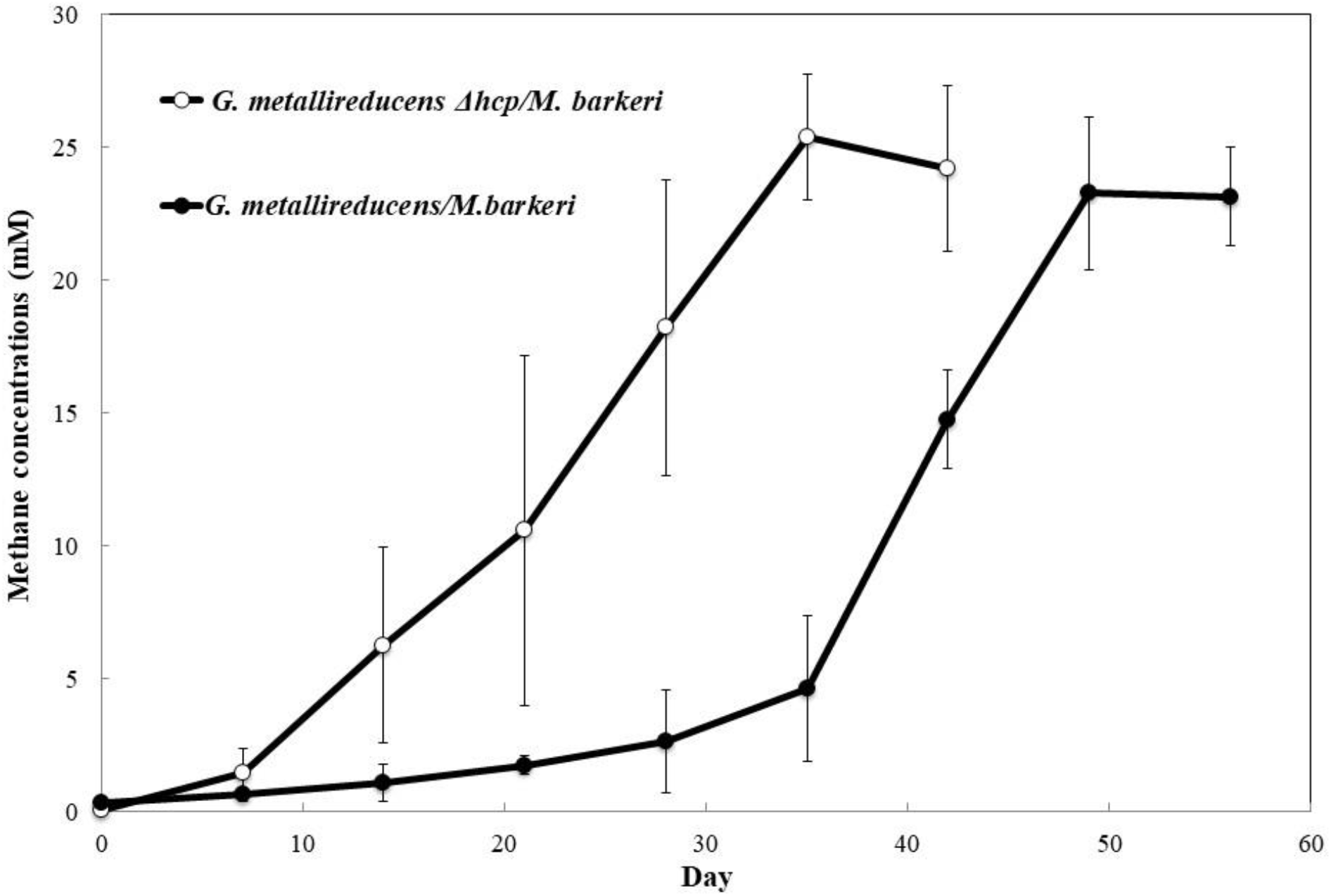
Methane production from DIET co-cultures initiated with the wild-type or *hcp*-deficient strain of *G. metallireducens* and wild-type *M. barkeri* during the first transfer. Each point and error bars represent the average and standard deviation of triplicate measurements.

### Disrupting the *hcp* gene in *G. metallireducens* has unexpected impact on extracellular electron transfer

The impact of the *hcp* deletion on extracellular electron transfer was evaluated. The Hcp-deficient strain grew faster than wild-type *G. metallireducens* when insoluble Fe(III) oxide was provided as the electron acceptor (Fig. 3). A complement strain containing the *hcp* gene *in trans* grew at rates similar to the wild-type strain, demonstrating that elimination of the T6SS impacted extracellular electron transfer capabilities.

**Figure 3.**
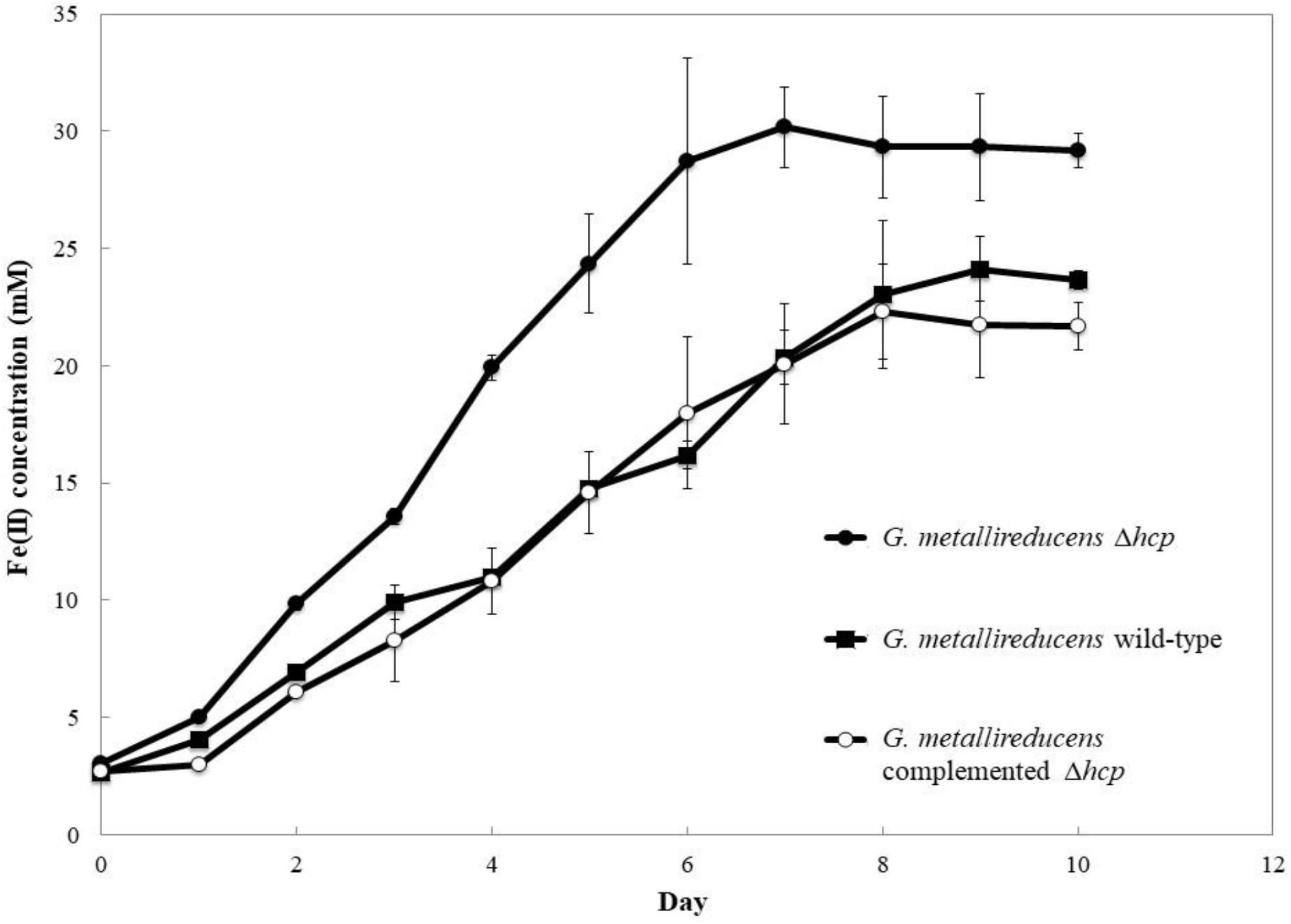
Fe(II) production by wild-type *G. metallireducens*, Δ*hcp*, and complemented Δ*hcp* strains during growth with acetate (20 mM) provided as the electron donor and Fe(III) oxide (50 mM) provided as the electron acceptor. Results and error bars represent triplicate cultures. The growth rate for Δ*hcp* was 1.5 (p-value=0.002) and 1.4 (p-value=0.02) times greater than wild-type or complemented Δ*hcp* strains.

Quantitative RT-PCR (Fig. 4) revealed that, compared to the wild-type strain, the Hcp-deficient strain of *G. metallireducens* more highly expressed genes for key outer-surface components previously shown to be important in extracellular electron transfer (Fig. 4; Table S6). These included genes for PilA, the monomer that is assembled into electrically conductive pili that are required for Fe(III) oxide reduction and DIET (25), as well as genes for the outer surface *c*-type cytochromes (Gmet_0930 and Gmet_0910) and a lipopolysaccharide protein (Gmet_2029) that are required for Fe(III) oxide reduction and expected to play an important role in DIET (23, 59, 80).

**Figure 4.**
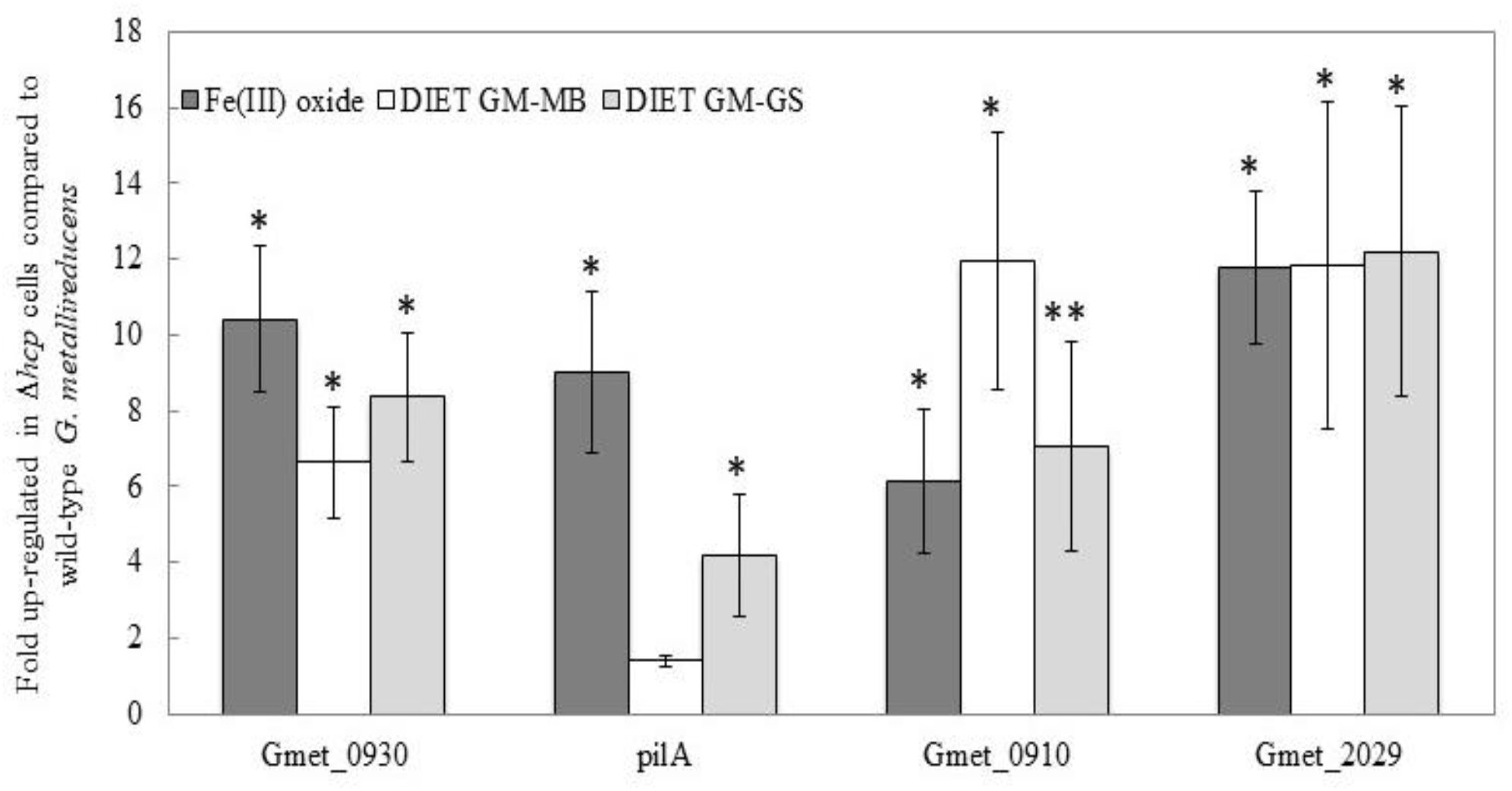
Results from quantitative RT-PCR using primers targeting various genes that code for proteins shown to be involved in extracellular electron transfer. mRNA used as template was extracted from cultures of *G. metallireducens* growing by Fe(III) respiration with acetate (20 mM) as the electron donor and Fe(III) oxide (50 mM) as the electron acceptor (Fe(III)-oxide), growing in co-culture with *M. barkeri* (DIET GM-MB), or growing in co-culture with *G. sulfurreducens* (DIET GM-GS). Results were calculated from triplicate biological and technical replicates using three different housekeeping genes as references (*proC, recA*, and *rpoB*). *represents p-values < 0.05; ** represents p-values <0.01. Further details regarding qRT-PCR results and p-values are available in Table S6.

Increased expression of these genes is a likely explanation for the accelerated Fe(III) oxide reduction and may also have contributed to accelerated DIET. The biosynthetic and energetic costs of deploying T6SS machinery is high (88). Therefore, it may be that eliminating some of this cost by deleting *hcp* enabled a greater investment in expression of outer-surface proteins important for extracellular electron transfer. Notably, T6SS genes were not up-regulated in *G. metallireducens/G. sulfurreducens* co-cultures that had undergone long-term adaptation to growth via DIET (23), suggesting that lowering expression of T6SS genes is part of *G. metallireducens*’ adaptive response to DIET-based growth.

### Implications

The results demonstrate that the expression of a T6SS can be detrimental for the establishment of DIET consortia and offer a new insight into the mechanisms by which conductive materials might facilitate DIET. It has previously been considered that conductive materials that are larger than cells, such as GAC, biochar, or carbon cloth accelerate the initiation of DIET because: 1) expression of electrically conductive pili and some outer-surface cytochromes is no longer necessary, conserving energy; and 2) it is easier for an electroactive microbe to establish electrical contact with a large conductive surface than small, disperse electrical contacts on another microbial cell (1, 89). The results presented here suggest that another benefit, in some instances, may be that conductive surfaces alleviate the need for close physical contact between DIET partners (60-62). Thus, DIET partners can ‘socially distance’ to avoid the possible negative impact of close physical associations as electrons zoom through the conductive material enabling the cells to connect remotely.

*G. metallireducens*’ high expression of its T6SS is clearly not in its best interest in the context of DIET in a defined laboratory co-culture, but *G. metallireducens* seems unlikely to exemplify the *Geobacter* species that participate in DIET in natural communities. *G. metallireducens* was recovered from an enrichment culture that selected for microbes rapidly growing via Fe(III) oxide reduction (69, 90), conditions likely to favor interspecies competition, not cooperation. Not all *Geobacter* species possess a T6SS (Table S1). Many other microbes that participate in DIET lack a T6SS, including *Prosthecochloris aestaurii* (16), *Syntrophus aciditrophicus*, (8), *Rhodoferax ferrireducens* (13), *Desulfovibrio* sp. JY (15), and *Rhodopseudomonas palustris* (18, 20) (Table S1b).

The finding that deletion of a major gene necessary for T6SS function in *G. metallireducens* unexpectedly increased expression of key components for extracellular electron transfer emphasizes a frequent problem in studies of *Geobacter* electromicrobiology. Adaption to gene deletions often result in changes in the mutant’s physiology beyond the direct function of the missing protein (65, 91, 92). Thus, multiple experimental approaches are warranted when developing models for *Geobacter* extracellular electron exchange.

