## Supplementary Materials for "Detrimental Impact of a Type VI Secretion System on Direct Interspecies Electron Transfer"

**Supplementary Figures**

**Supplementary Figure S1.** Phylogenetic tree displaying a comparison of *Geobacter* TssC proteins to sequences from other Gram-negative bacteria

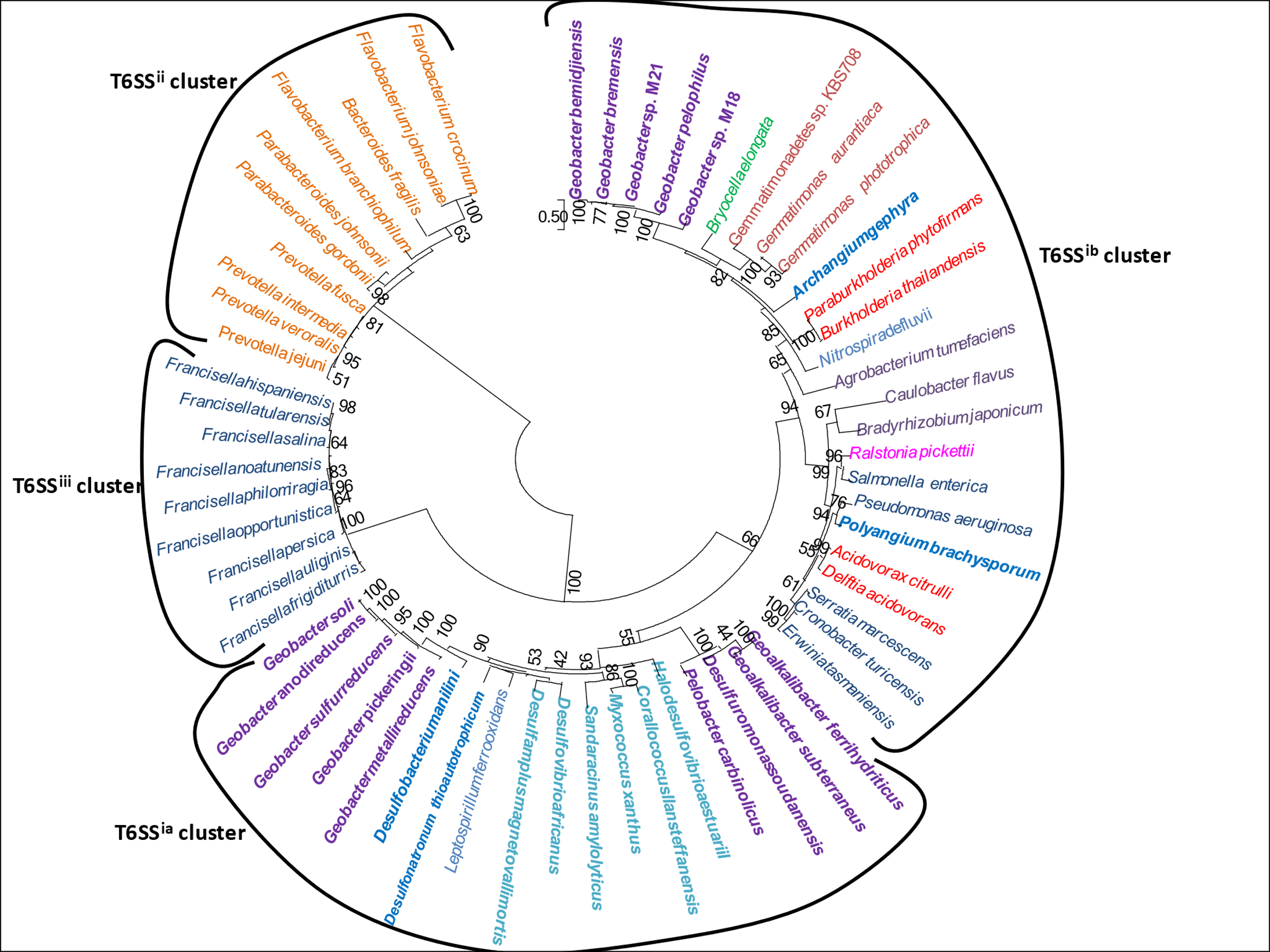

**Supplementary Figure S2.** Multidimensional scaling (MDS) plot showing distances between samples based on the biological coefficient of variation (BCV) method.

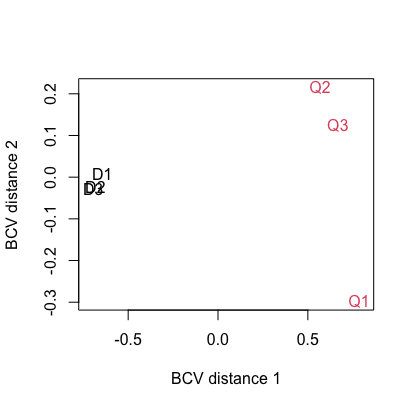

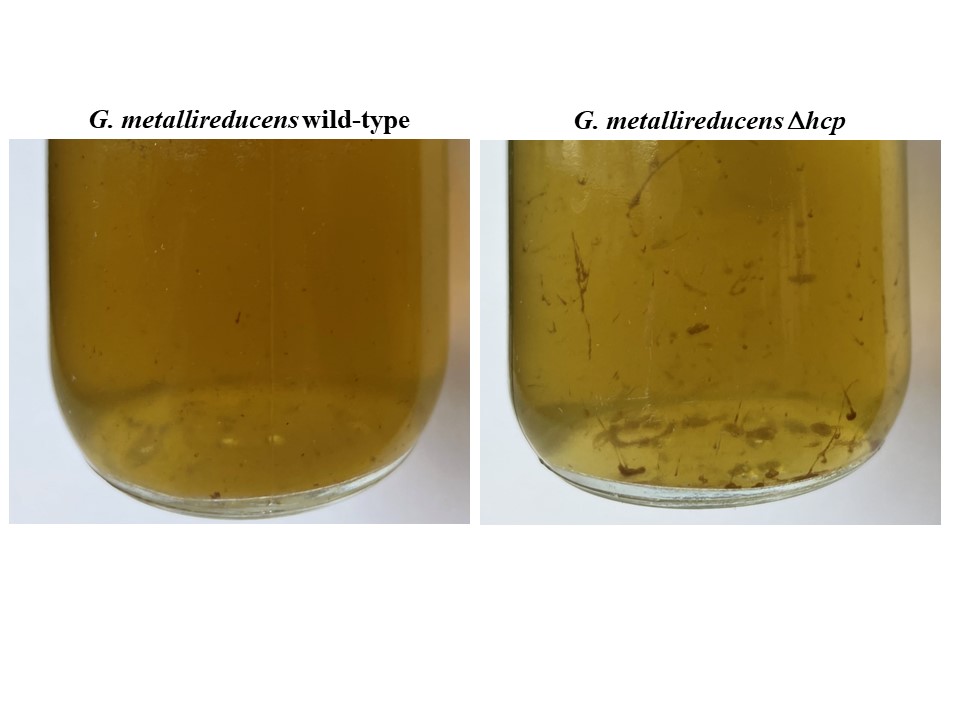
**Supplementary Figure S3**. Images of *G. metallireducens* wild-type and *G. metallireducens* Δ*hcp* grown in medium with Fe(III) citrate as the electron acceptor when Fe(II) concentrations reached approximately 40mM

**Supplementary Tables**

**Supplementary Table S3.** Primers used for qRT-PCR analysis of *G. metallireducens* strains grown under various culturing conditions.

| **Primer names** | **forward primer sequence (5'to3')** | **reverse primer sequence (5'to3')** |
| --- | --- | --- |
| Gmet_recA-446f/549r | TCGTTCCCAAGGCCGAGATC | GGAGATGATGCCGGTGAGCT |
| Gmet_rpoB-1483f/1587r | CTGATGCCCCACGACCTGAT | AAGCGGGTTTGTCTGGTCCA |
| Gmet_proC-224f/352r | AGGTATTTCGGGGCATGGCT | GGATCACCCGGGAAGAGTCG |
| Gmet_pilA-34f/150r | ACCCTCATCGAGCTGCTGAT | CAAGTCAGACTCGGCAGCAG |
| Gmet_0930-1197f/1296r | ACTTCCGCTCCTCACCATCA | GGTGTTCACGTCGGTAGCAG |
| Gmet_omcF-1062f/1182r | CATCACATCAGCACCGCACT | CAGGATAGTGTGAGCCCCGA |
| Gmet_2029-472f/615r | AAGCGGCAGGATTCCTTTGG | GAGAAGCGCCTCGTCAATCC |
| Gmet_2839-189f/317r | GATCACCTGCTTCGACTGCC | GGATTCTTGTGGCACCCGAG |

**Supplementary Table S4. Primers used for mutant construction.**

| **Primer Name** | **Purpose** | **Sequence (5’ to 3’)** |
| --- | --- | --- |
| 0280_up | GS15 Δ0280::Sp^r^ | CATCGCCCTCACCATGAGAAAGG |
| 0280_AvrII_up | GS15 Δ0280::Sp^r^ | CTGCCTAGGCCACAAGGAGGAAGTAATCAATG^#^ |
| 0280_AvrII_dn | GS15 Δ0280::Sp^r^ | CAGCCTAGGCAACGGCATCGAAGCCGAG^#^ |
| 0280_dn | GS15 Δ0280::Sp^r^ | ATAGCGGATGCTTCCGACAG |
| 3174_up | PCA Δ3174::Sp^r^ | CCGCGAACTCGGTCCAGAA |
| 3174_AvrII_up | PCA Δ3174::Sp^r^ | CTGCCTAGGCATCTGCTGCTACCTCCTTG^#^ |
| 3174_AvrII_dn | PCA Δ3174::Sp^r^ | TAGCCTAGGCAACGGTATCGAGGCTGAGGATT^#^ |
| 3174_dn | PCA Δ3174::Sp^r^ | GAAGTATGTTACGCCATGTTCG |
| 0280_comp_fwd | GS15 0280 complementation | GTATCTAGACACAAGGAGGAAGTAATCAATG^##^ |
| 0280_comp_rev | GS15 0280 complementation | GTAGGTACCAGGAACATCGGACAACTAACTA^###^ |

^#^*AvrII* restriction site is underlined

^##^XbaI restriction site is underlined

^###^KpnI restriction site it underlined

**Supplementary Table 5.** Differences in transcript abundance for all 13 core T6SS genes in *G. sulfurreducens* grown via DIET compared to QUIET and for genes coding for putative T6SS effector and associated immunity proteins (P-value cutoff ≤ 0.05).

ND: no difference

tse: T6SS effector protein; tsi: T6SS immunity protein

| Gene | Name | Annotation | Fold-change  DIET vs. QUIET | p-value |
| --- | --- | --- | --- | --- |
| **Core T6SS Genes** | | | | |
| GSU3165 | *tssL* | membrane core complex protein | ND |  |
| GSU3166 | *tssM* | membrane core complex protein | 2.81 | 0.03 |
| GSU3167 | *tssA* | baseplate complex protein | 250.65 | 3.90x10^-5^ |
| GSU3172 | *tssB* | Sheath protein | ND | ND |
| GSU3173 | *tssC* | Sheath protein | 23.07 | 3.80x10^-7^ |
| GSU3174 | *hcp* | inner tube protein | 11.76 | 0.0003 |
| GSU3177 | *vrgG* | puncturing device | ND | ND |
| GSU0428 | *tssJ* | membrane core complex protein | ND | ND |
| GSU0429 | *tssK* | baseplate complex protein | 4.80 | 0.0003 |
| GSU0430 | *tssE* | baseplate complex protein | ND | ND |
| GSU0431 | *tssF* | baseplate complex protein | ND | ND |
| GSU0432 | *tssG* | baseplate complex protein | ND | ND |
| GSU0433 | *tssH* | ClpV1 protease involved in sheath recycling | 4.33 | 0.007 |
| **Effector/ Immunity Genes** | | | | |
| GSU3168 | *tse1* | fatty acid metabolism protein | ND |  |
| GSU3169 |  | PAAR-like DUF4150 domain protein | ND |  |
| GSU3171 | *tse2* | PGAP1 domain protein; phospholipase | ND |  |
| GSU3176 | *tse3* | lysM domain protein | ND |  |
| GSU3181 | *tse4* | fatty acid metabolism protein | 51.42 | 0.01 |
| GSU3182 |  | PAAR-like DUF4150 domain protein | ND |  |
| GSU3182 |  | PAAR-like DUF4150 domain protein | ND |  |

**Supplementary Table S6.** Results from quantitative RT-PCR using the housekeeping genes, *recA*, *proC*, and *rpoB* as standards. Values represent fold up-regulated in Hcp-deficient (∆Gmet_0280) *G. metallireducens* strains compared to wild-type strains grown with acetate (20 mM) as the electron donor and Fe(III) oxide (50 mM) as the electron acceptor (Fe(III)-oxide), grown in co-culture with *M. barkeri* with ethanol (20 mM) as the electron donor and CO_2_ as the electron acceptor (GM-MB), or grown in co-culture with *G. sulfurreducens* with ethanol (20 mM) as the electron donor and fumarate (45 mM) as the electron acceptor. All values were obtained from triplicate biological and technical replicates and p-values were calculated by ANOVA using the R statistical package.

|  | **Gmet_0930/*recA* fold difference** | **p-value** | **Gmet_0930/*proC* fold difference** | **p-value** | **Gmet_0930/ *rpoB* fold difference** | **p-value** | **average fold difference** | **standard deviation** | **average**  **p-value** |
| --- | --- | --- | --- | --- | --- | --- | --- | --- | --- |
| Fe(III) oxide | 10.76 | 0.04 | 8.33 | 0.01 | 12.13 | 0.035 | 10.41 | 1.92 | 0.031 |
| DIET GM-MB | 5.21 | 0.01 | 6.59 | 0.05 | 8.12 | 0.016 | 6.64 | 1.46 | 0.024 |
| DIET GM-GS | 6.74 | 0.05 | 8.22 | 0.009 | 10.14 | 0.02 | 8.37 | 1.71 | 0.025 |
|  | ***pilA*/*recA* fold difference** |  | ***pilA*/*proC* fold difference** |  | ***pilA*/*rpoB* fold difference** |  | **average fold difference** | **standard deviation** | **average**  **p-value** |
| Fe(III) oxide | 8.11 | 0.05 | 7.63 | 0.03 | 11.32 | 0.006 | 9.02 | 2.01 | 0.028 |
| DIET GM-MB | 1.57 | 0.20 | 1.27 | 0.39 | 1.4 | 0.40 | 1.41 | 0.15 | 0.331 |
| DIET GM-GS | 2.48 | 0.05 | 5.72 | 0.03 | 4.33 | 0.004 | 4.18 | 1.63 | 0.028 |
|  | **Gmet_0910/*recA* fold difference** |  | **Gmet_0910/*proC*** |  | **Gmet_0910/*rpoB* fold difference** |  | **average fold difference** | **standard deviation** | **average**  **p-value** |
| Fe(III) oxide | 4.0 | 0.05 | 8.29 | 0.02 | 6.08 | 0.002 | 6.12 | 2.15 | 0.024 |
| DIET GM-MB | 8.24 | 0.02 | 14.96 | 0.02 | 12.58 | 0.02 | 11.93 | 3.41 | 0.024 |
| DIET GM-GS | 4.2 | 0.007 | 9.69 | 0.01 | 7.33 | 0.01 | 7.07 | 2.75 | 0.009 |
|  | **Gmet_2839/*recA* fold difference** |  | **Gmet_2839/*proC*** |  | **Gmet_2839/*rpoB* fold difference** |  | **average fold difference** | **standard deviation** | **average**  **p-value** |
| Fe(III) oxide | 1.5 | 0.71 | 1.17 | 0.76 | 2.28 | 0.15 | 1.65 | 0.57 | 0.541 |
| DIET GM-MB | 1.23 | 0.19 | 1.1 | 0.72 | 1.1 | 0.24 | 1.14 | 0.08 | 0.382 |
| DIET GM-GS | 1.1 | 0.94 | 2.09 | 0.55 | 1.58 | 0.69 | 1.59 | 0.50 | 0.725 |
|  | **Gmet_2029/*recA* fold difference** |  | **Gmet_2029/*proC* fold difference** |  | **Gmet_2029/*rpoB* fold difference** |  | **average fold difference** | **standard deviation** | **average**  **p-value** |
| Fe(III) oxide | 13.64 | 0.05 | 12.08 | 0.06 | 9.62 | 0.004 | 11.78 | 2.03 | 0.038 |
| DIET GM-MB | 10.32 | 0.02 | 8.44 | 0.04 | 16.69 | 0.01 | 11.82 | 4.32 | 0.024 |
| DIET GM-GS | 8.15 | 0.02 | 15.81 | 0.01 | 12.64 | 0.05 | 12.20 | 3.85 | 0.030 |

**Supplemental Text**

**Phylogeny of *Geobacter* T6SSs**

The type VI secretion system (T6SS) is composed of 13 core proteins that collectively form a contractile nanomachine that injects toxins into target cells in a manner similar to bacteriophage (1). This nanomachine is composed of four main parts; the baseplate, the membrane complex, the sheath, and the puncturing device. The baseplate complex serves as a platform for contractile tail elongation and it is composed of four subunits (TssE, TssF, TssG, TssK). The membrane core complex serves as the T6SS docking station and platform for baseplate assembly and it is composed of three proteins (TssJ, TssL, TssM). The sheath is composed of TssB and TssC and the inner tube consists stacked hexamers of the Hcp protein. TssA properly attaches the sheath to the baseplate and helps stabilize sheath structure, while TssH is a ClpV protease that helps recycle the sheath. The puncturing device located at the top of the inner tube is composed of a VgrG trimer. In some cases, an additional Pro-Ala-Ala-Ala (PAAR)- repeat protein is found in the operon that helps sharpen the tip.

Effector molecules are loaded onto the tip of the VgrG trimer and are released into the target cell once the cell’s membrane has been pierced. While there is much conservation among the 13 proteins comprising the actual T6SS structure, there is a considerable amount of variation between effector proteins. Many of the effector proteins that have been studied to date are injected directly into cells and target bacterial cell membranes, cell walls, or nucleic acids (2-4). However, extracellular effectors that facilitate metal uptake (5-7) have also been identified.

A genome-wide comparison of currently sequenced *Geobacter* genomes revealed that 10 out of the 16 sequenced *Geobacter* species have genes coding for the necessary components of T6SSs. These species include: *G. metallireducens*, *G. sulfurreducens*, *G. pickeringii*, *G. soli*, *G. anodireducens*, *G. bemidjiensis*, *G. bremensis*, strain M18, strain M21, and *G. pelophilus* (Table S1). Several other closely related genera from the order *Desulfuromondales* also appear to have T6SSs; *Desulfuromonas soudanensis*, *Geoalkalibacter ferrihydriticus*, *Geoalkalibacter subterraneus*, and *Pelobacter carbinolicus* (Table S1). In 5 of these genomes, the T6SS genes are separated into two main clusters, while the T6SS genes are found in a single operon in the other *Geobacter* genomes (Table 1; Table S1).

Two different types of effectors have been identified in T6SSs. Specialized effectors are fused to the C-terminus of T6SS VgrG, Hcp or PAAR-domain containing proteins, while cargo effectors need help from a chaperone or adaptor protein to be loaded onto the Hcp tube or the VgrG spike (4). It appears that *Geobacter* and other *Desulfuromondales* species utilize cargo effector mechanisms, because chaperone-like proteins were identified in all T6SS gene clusters (Table S1). Similar to the *Agrobacterium tumefaciens* T6SS, all *Geobacter* species with T6SS genes divided into two gene clusters and other *Desulfuromondales* species have genes coding for putative DUF2169-containing chaperone proteins and genes for PAAR-like DUF4150 proteins (4, 8). The PAAR-like DUF4150 domain protein is thought to form a sharp conical extension on the VgrG spike and to help attach effectors to VgrG (9). The DUF2169 containing gene, frequently found upstream from DUF4150, codes for a chaperone required for effector-dependent antibacterial activity by *A. tumefaciens* (8). All of the *Geobacter* with T6SS genes concentrated within one region of the genome lacked DUF4150- and DUF2169-containing genes, and instead had a gene encoding another DUF4123-containing chaperone required for effector translocation to VgrG in *Vibrio cholerae* (4, 10, 11).

Comparison of *G. metallireducens* and *G. sulfurreducens* T6SS operons to other well-characterized T6SSs identified several genes that could potentially encode effectors and their associated immunity proteins which are needed to ensure that the host cell is not destroyed by its own toxins. In the *G. metallireducens* genome, a gene coding for a D-alanyl-D-alanine carboxypeptidase with a peptidoglycan-binding site (Gmet_0285) is located next to the VgrG puncture protein. Another gene that could potentially serve as an immunity protein is Gmet_0284 which has an NTF2-like domain that is found in immunity proteins from various bacterial polymorphic toxin systems (12, 13). The operon also has a putative fatty acid degradation gene (Gmet_0287) located next to a PAAR-like DUF4150 domain protein (Gmet_0288), similar to a T6SS gene cluster found in *Pseudomonas aeruginosa* (PA0082-PA0101) and a gene coding for a putative phospholipase effector protein (Gmet_0290) located next to a gene coding for a putative chaperone protein with a DUF2169 domain.

Similar to *G. metallireducens*, one of the T6SS gene clusters in *G. sulfurreducens* also has several genes coding for putative effectors and their associated immunity and chaperone proteins. GSU3168 and GSU3181 both code for a putative lipase located next to a gene coding for a PAAR-like DUF4150 domain protein; GSU3171 codes for a putative phospholipase; and GSU3176 which is located adjacent to the gene for the VgrG spike encodes a protein with a lysM domain that could be involved in cell wall degradation. In addition, GSU3184 encodes a putative amidohydrolase protein located in the same vicinity as a gene coding for a putative DUF2169 chaperone protein (GSU3186).

Many pathogenic Gram-negative bacteria appear to have multiple copies of T6SS gene clusters. For example, *Proteus mirabilis* has five T6SS gene clusters (14), *Burkholderia mallei* and *Salmonella enterica* have at least four T6SS gene clusters (15, 16), and *Pseudomonas aeruginosa* has at least three functional T6SS gene clusters (17). Studies have shown that each of these T6SS clusters encode structures with different functions (i.e. pathogenicity, stress response, intraspecific cooperativity) (18). It does not appear that *Geobacter* and other deltaproteobacteria analyzed in this study have more than one set of T6SS genes.

The majority of T6SS proteins that have been characterized to date appear to fall into three different phylogenetic groups (19-23), with the exception of *Ca. Amoebophilus asiaticus* which has been proposed to form its own distinct phylogenetic group, T6SS^iv^ (24). Previous studies have found that T6SS^i^ includes sequences from Proteobacteria, *Francisella* sequences form the T6SS^ii^ clade, and T6SS^iii^ is composed of *Bacteroidetes* T6SS sequences. These phylogenies compared the large sheath protein (TssC) (19-21), the small sheath protein (TssA) (22), and the baseplate protein TssF (23). Comparison of *Geobacter* TssC proteins to sequences from other Gram-negative bacteria showed that *Geobacter* T6SS clusters within the T6SS^i^ clade (Figure S1). However, *Geobacter* T6SS proteins can be further divided into 2 subclades (T6SS^ia^ and T6SS^ib^). While TssC from those *Geobacter* with their T6SS genes concentrated in a single operon (*G. bremensis*, *G. bemidjiensis*, *G. pickeringii*, strain M21, strain M18) clustered within the T6SS^ia^ clade, TssC from *Geobacter* with T6SS genes divided into two regions within the genome (*G. sulfurreducens*, *G. metallireducens*, *G. soli*, *G. anodireducens*, *G. pelophilus*) fell within the T6SS^ib^ clade.
